# Surface-engineered extracellular vesicles to modulate antigen-specific T cell expansion for cancer immunotherapy

**DOI:** 10.1101/2023.09.25.559260

**Authors:** Xiabing Lyu, Tomoyoshi Yamano, Shota Imai, Toan Van Le, Dilireba Bolidong, Makie Ueda, Shota Warashina, Hidefumi Mukai, Seigo Hayashi, Kazutaka Matoba, Taito Nishino, Rikinari Hanayama

## Abstract

Extracellular vesicles (EVs), including exosomes, are emerging as novel mediators of cell-cell communications, involved in various processes such as immune activation and immunosuppression. Despite the recent development of several EVs-based cancer immunotherapies, their clinical efficacy remained limited. Here, using fusion with tetraspanin as one of the EV engineering techniques, we created antigen-presenting extracellular vesicles (AP-EVs) to reproduce the functional characteristics of professional antigen-presenting cells (APCs). AP-EVs were also equipped with surface-bound IL-2, a feature not inherent to APCs, which facilitated selective delivery of IL-2 to antigen-specific CD8^+^ T cells. AP-EVs were engineered to express a peptide-major histocompatibility class I (pMHCI) complex, a costimulatory CD80 molecule, and IL-2, allowing the simultaneous presentation of multiple immune modulators to antigen-specific CD8^+^ T cells. This promoted the clonal expansion and differentiation of antigen-specific cytotoxic T lymphocytes, leading to potent anticancer immune responses. Combination therapy with AP-EVs and anti-PD-1 demonstrated enhanced anticancer immunity against established tumors compared with anti-PD-1 monotherapy. Our engineered EVs represent a novel effective strategy for cancer immunotherapy.

## Introduction

Extracellular vesicles (EVs), including exosomes, are secreted by most cells. Extracellular vesicles contain various proteins, lipids, and RNAs that induce functional and physiological changes in target cells^1^. They are considered promising nano-vesicles in clinical settings owing to their biocompatibility, low immunogenicity, and high efficacy for drug delivery^2^. Recent studies have demonstrated that EVs released by cancer cells or antigen-presenting cells (APCs) modulate immune responses^3^. For example, cancer-derived EVs express PD-L1, a ligand of the inhibitory receptor PD-1 on T cells, which hampers the immune response against cancer^4,5^. In contrast, mature dendritic cell (DC)-derived exosomes express peptide-major histocompatibility complex (MHC), molecules that can directly or indirectly stimulate T cells^6^. Previous studies have reported cancer immunotherapies using DC-derived exosomes by exploiting their stimulatory properties^7–9^. However, only a few clinical studies have demonstrated a therapeutic effect on cancer progression, possibly because of the limitations of costimulatory molecules and cytokines^10^.

Cytokines play a pivotal role in regulating immune homeostasis, and their quantities are crucial in clinical immunotherapy^11,12^. Cytokines are highly effective locally but have severe adverse effects when used systemically^13^. For example, IL-2 has been clinically used to improve T cell immunity in patients with metastatic melanoma and renal cell carcinoma^13,14^. However, a high dose of IL-2 activates endothelial cells, which express high-affinity IL-2Rα, leading to vascular leak syndrome^15^. In addition, IL-2 increases the number of regulatory T cells (Tregs), weakening the immune responses to cancer^16^. Several strategies have been developed to overcome these challenges. For example, some studies have reported the engineering of IL-2 variants that selectively stimulate the memory phenotype CD8^+^ T and natural killer (NK) cells. These IL-2 variants have been engineered to bind preferentially to IL-2Rβ, thereby reducing the undesirable expansion of Tregs and associated adverse events^17–19^. Another approach involves the creation of a fusion protein that combines IL-2 with the Orthopoxvirus MHC class I-like protein (OMCP), a known ligand of NKG2D, directing IL-2 specifically to NKG2D-positive cells^20^. However, despite these advances in IL-2 variants and IL-2-OMCP fusion proteins, IL-2 variants and IL-2-OMCP fusion protein primarily act on polyclonal CD8^+^ T cells, many of which lack antigen specificity against tumors. Thus, the targeted delivery of IL-2 to antigen-specific CD8^+^ T cells remains challenging.

Given the importance of targeted cytokine delivery and the potential of EVs in immune modulation, there is a notable demand to develop strategies that harness the advantages of both. This study aims to engineer antigen-presenting extracellular vesicles (AP-EVs) to bridge this gap and explore their potential in enhancing specific T cell responses.

## Results

### Simultaneous expression of immune modulators on EVs

To display the protein of interest (POI) on the surfaces of EVs, we employed a technique of fusion with tetraspanins, such as CD9 or CD81 (Fig. 1a)^21^. First, we designed the CD81 fusion protein with a single-chain MHCI trimer (scMHCI) consisting of an OVA peptide, β2m, and H2-K^b^ molecule (Extended Data Fig.1a)^22^. We integrated IL-2 into the second extracellular loop of CD81 via a GS linker to ensure its robust and functional display on the EVs (Extended Data Fig.1b). Concurrently, CD80 was fused with CD9 (Extended Data Fig.1c). Subsequently, human embryonic kidney (HEK293) cells that stably expressed scMHCI-CD81-IL-2 and CD80-CD9 were used to harvest EVs from their culture supernatant. High expression levels of scMHCI, CD80, and IL-2 present on the EVs from HEK293 cells were observed through flow cytometry (Fig.1b). We named these EVs antigen presenting extracellular vesicles (AP-EVs). To verify the concurrent expression of CD9 and CD81 fusion proteins on individual EVs, HEK293T cells were co-transfected with CD9-GFP and CD81-RFP constructs and EVs were isolated from the culture supernatant. In line with prior observations^23^, confocal microscopy analyses revealed that approximately 80% of the EVs co-expressed GFP and RFP (Fig.1c, Extended Data Fig.1d), suggesting that a majority of AP-EVs simultaneously display scMHCI-CD81-IL-2 and CD80-CD9 on a single EV. Nanoparticle tracking analysis (NTA) showed that the average size of AP-EVs is approximately 150nm, which is comparable to that of non-modified control EVs (Fig.1d), and atomic force microscopy (AFM) showed that AP-EVs have morphological and physical properties similar to those of EVs (Fig.1e, f, Extended Data Fig.1e) ^24^.

**Figure 1.**
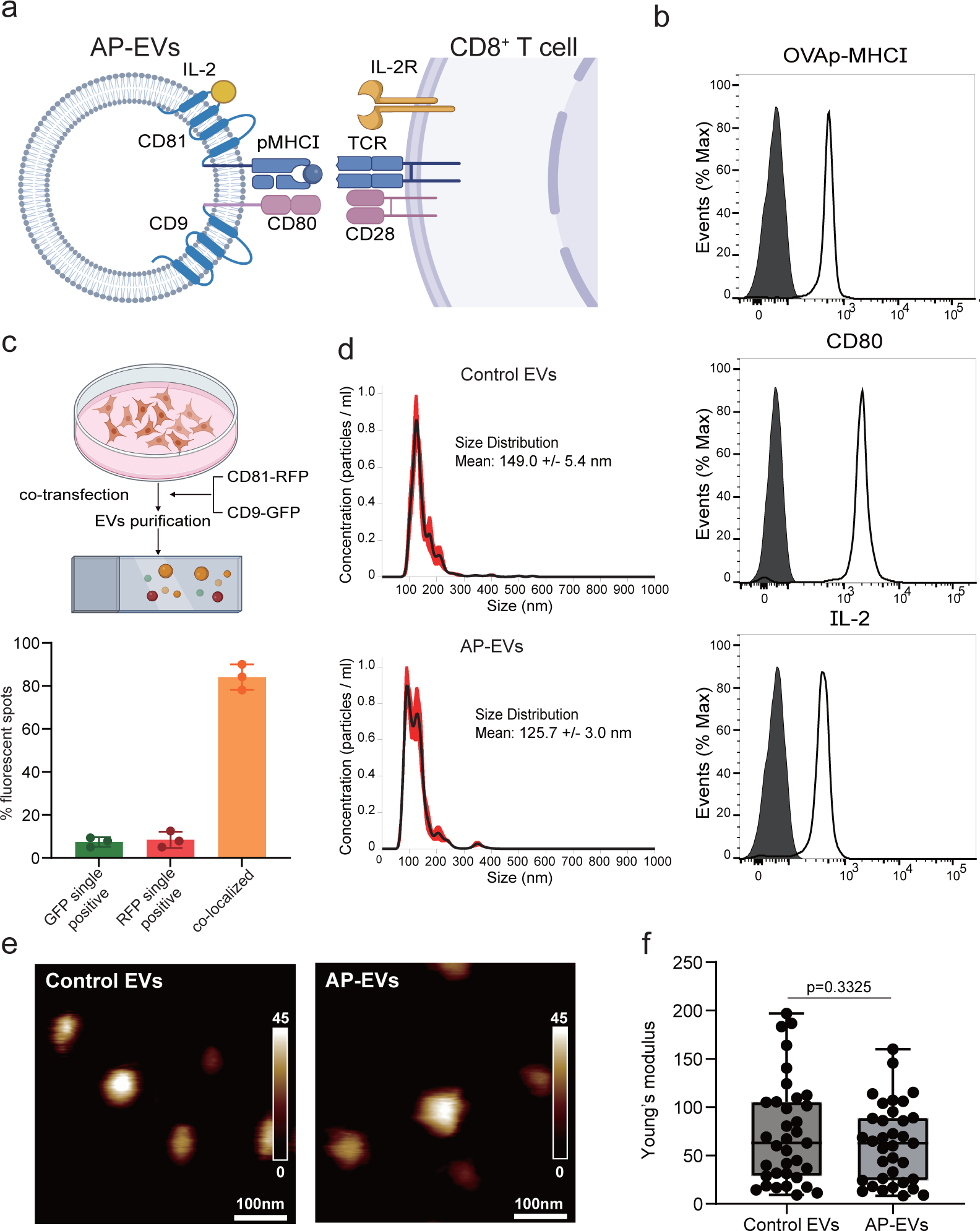
Establishment of antigen-presenting EVs (AP-EVs). (a) Schematic representation of AP-EVs. Functional pMHCI complex, CD80, and IL-2 interact with TCR, CD28, and IL-2R, respectively, on antigen-specific CD8^+^ T cells. (b) Surface expression of OVAp-MHCI complex, CD80, and IL-2 on EVs was assessed using flow cytometry. Filled greys represent control EVs, and open blacks represent AP-EVs. (c) HEK293 T cells were co-transfected with CD81-RFP and CD9-GFP, and the EVs were isolated. These isolated EVs were placed onto glass slides and observed under confocal microscopy. Colocalization of RFP and GFP was quantified. (d) Size distribution between the control EVs and AP-EVs (Red shaded region = standard error of the mean. (e) Control EVs and AP-EVs showed similar morphology under high-speed atomic force microscopy (HS-AFM). (f) Estimation of the elastic modulus (Young’s modulus) for EVs and AP-EVs were performed. Data (b-f) are representative of two independent experiments.

### Selective expansion of antigen-specific CD8^+^ T cells with AP-EVs

Previous reports have shown that DC-derived exosomes can directly prime effector T cells but are ineffective in stimulating naïve T cells^25,26^. We investigated if AP-EVs can stimulate naïve CD8^+^ T cells; therefore, OVA-specific OT-I CD8^+^ T cells labeled with Cell Trace Violet (CTV) were cultured with various doses of AP-EVs or control EVs. After three days of cultivation, a dose-dependent expansion of OT-I T cells by AP-EVs was observed following treatment with AP-EVs: expansion was over 60% at 3 µg/ml, above 40% at 1 µg/ml, just under 20% at 0.3 µg/ml, and minimal at 0.1 µg/ml (Fig.2a, Extended Data Fig.2), indicating that AP-EVs directly activated naïve CD8^+^ T cells. Subsequently, we examined which components (signal1: scMHCI, signal2: CD80, signal3: IL-2) on AP-EVs were minimally required for T cell activation. We prepared EVs expressing signal1 alone, signal1 and 2, signal1 and 3, or signal1, 2, and 3 (Fig.2b), and cocultured them with OT-I T cells. The EVs that expresing signal 1 alone did not efficiently activate OT-I T cells (Fig. 2c). In contrast, co-expression with signals 2 or 3 strongly stimulated the proliferation of OT-I T cells, and the co-expression of signal1, 2, and 3 showed the maximum effect. This indicated that signals 2 and 3 on AP-EVs synergistically enhanced CD8^+^ T cell expansion. Moreover, OT-I T cells activated by AP-EVs produced IFN-γ and granzyme B, suggesting the conversion of naïve T cells into cytotoxic T lymphocytes (CTLs) (Fig. 2d).

**Figure 2.**
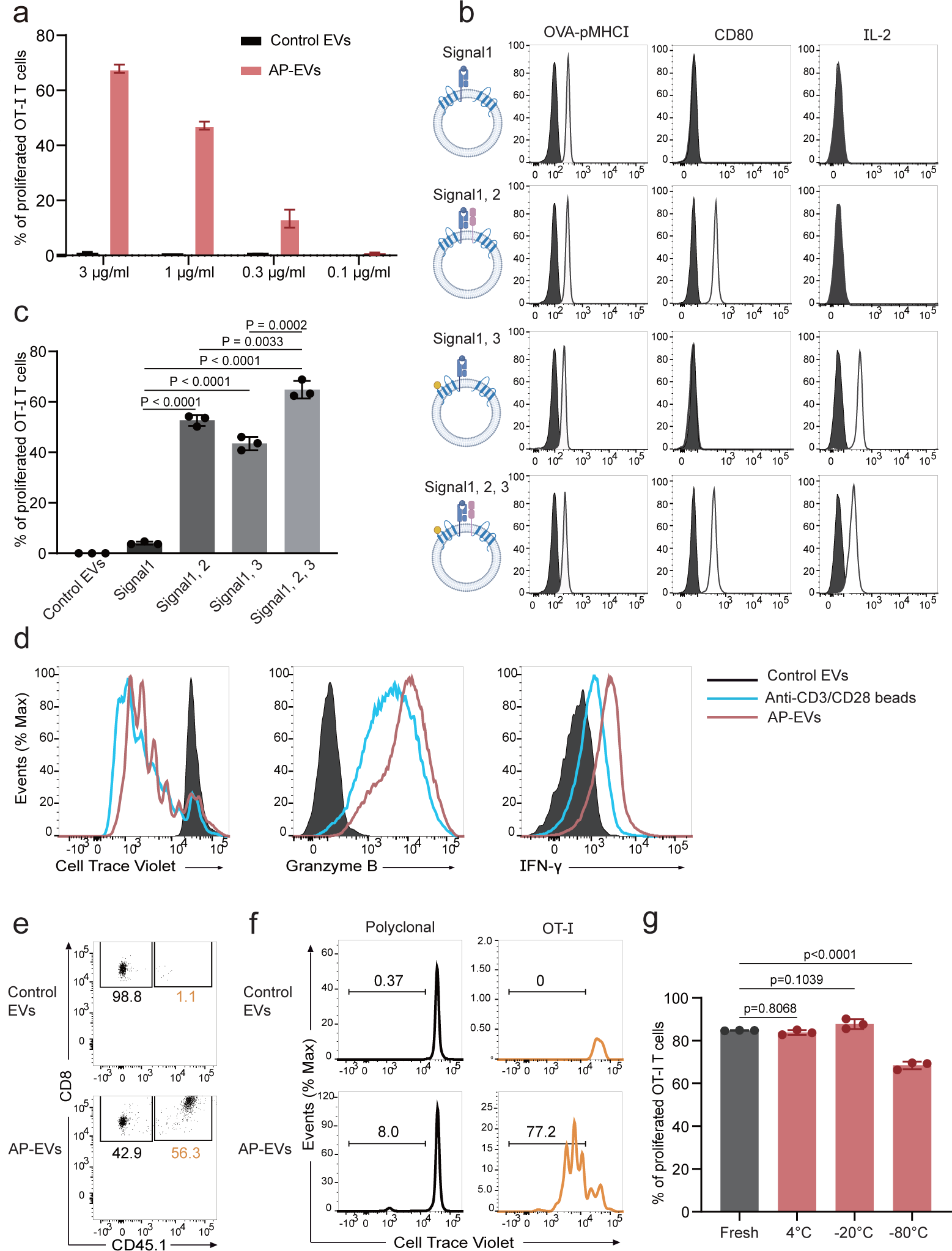
Direct activation of antigen-specific CD8^+^ T cells by AP-EVs. (a) Bar graph representing the percentage of proliferated OT-I T cells under different concentrations of EVs (black) and AP-EVs (red). (b) Surface expression of the OVAp-MHCI complex, CD80, and IL-2 on EVs through different signals (alone and in combination). (c) The stimulatory capability of EVs expressing individual signals (1, 2, and 3). CTV-labeled OT-I T cells were cultured with each type of EV and the proliferation of OT-I T cells was quantified using flow cytometry. P-values were calculated using a one-way analysis of variance (ANOVA) followed by the Dunnett test. (d) The productions of granzyme B and IFN-γ by CTV-labeled OT-I T cells cultured with AP-EVs were analyzed by intracellular staining. (e) Flow cytometric plots of CTV-labeled OT-I T cells (CD45.1) and polyclonal T cells (CD45.2) that was cultured with either AP-EVs or control EV. (f) Proliferation of OT-I and polyclonal T cells after culturing with control EVs or AP-EVs was determined by flow cytometry. (g) OT-I proliferation after culturing with AP-EVs stored at 4°C, −20°C, and −80° for three months or freshly prepared AP-EVs. P-values were determined using a one-way analysis of variance (ANOVA) followed by the Dunnett test. The proliferation of OT-I T cells was measured by flow cytometry. Data (a, b, c, e, f, g) are representative of two independent experiments.

OT-I T cells were cocultured with polyclonal CD8^+^ T cells at a 5:95 ratio to determine whether AP-EVs could specifically stimulate CD8^+^ T cells. Further, control EVs or AP-EVs were added to the culture. While control EVs maintained the initial ratio of OT-I to polyclonal CD8^+^ T cells, AP-EVs selectively expanded the OT-I T cell population in vitro (Fig. 2e, f). Although we utilized freshly isolated AP-EVs in this study, we observed that AP-EVs stored for up to 3 months at 4°C or −20°C (but not at −80°C) demonstrated activity comparable to that of fresh EVs (Fig. 2f). Collectively, these results indicate that AP-EVs, which directly present an antigen, a costimulatory molecule, and a cytokine, induce robust proliferation of antigen-specific CD8^+^ T cells.

### In vivo expansion of antigen-specific CD8^+^ T cells by AP-EVs

We next sought to determine if AP-EVs could selectively stimulate antigen-specific CD8^+^ T cells in vivo without causing severe adverse effects. This is important because using IL-2/anti-IL-2 mAb for in vivo CD8^+^ T cell expansion often leads to nonspecific activation and side effects such as splenomegaly^17^. To test this, we transferred a mixture of congenically marked OT-I T cells and polyclonal T cells (in a 1:1 ratio) to recipient CD45.1^+^CD45.2^+^ mice, followed by intravenous injection of AP-EVs, control EVs, or IL-2/anti-IL-2 mAb. After 4 days, donor T cells were analyzed (Fig.3a). Consistent with the previous finding^27^, the IL-2/anti-IL-2 mAb expanded both OT-I and polyclonal T cells. In contrast, AP-EVs exhibited selective expansion of OT-I T cells in the spleen (Fig.3a, b, c) and lymph nodes (LNs) (Extended Data Fig.3a, b, c), with the majority of OT-I T cells acquiring the CD44^hi^CD62L^low^ effector memory or effector phenotype (Fig. 3d, Extended Data Fig.3d). Additionally, a subset of OT-I T cells differentiated into IFN-γ positive CTLs (Fig.3e, Extended Data Fig.3e). Next, we evaluated the toxicity of AP-EVs. Their administration did not present any apparent adverse effects; no marked changes were observed in body weight, histological examinations revealed no tissue abnormalities, and blood chemistry analysis results were consistent with the vehicle-treated controls (Extended Data Fig.4a, b, c, d). Additionally, no expansion of polyclonal CD4^+^ T cells, polyclonal CD8^+^ T cells, NK cells, or Tregs was observed (Extended Data Fig.5a, b, c, d).

**Figure 3.**
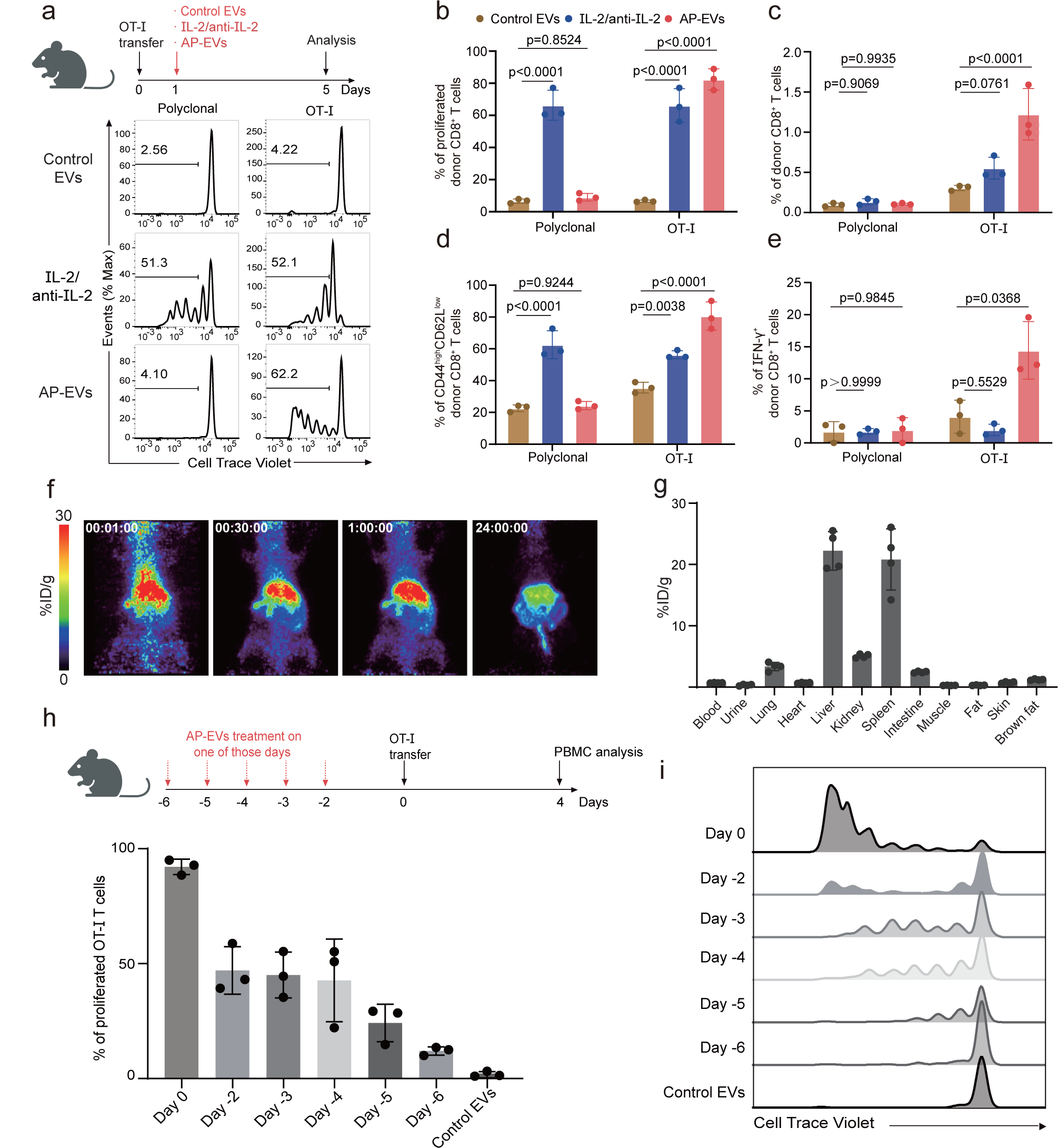
In vivo activation of antigen-specific CD8^+^ T cells by AP-EVs. (a) CTV-labeled OT-I (CD45.1) and polyclonal T cells (CD45.2) were transferred into CD45.1^+^CD45.2^+^ recipient mice. The following day, mice were administered either control EVs, IL-2/anti-IL-2 mAb, or AP-EVs. The activation status was analyzed 5 days later (n=3 mice per group). (b) The donor CD8^+^ T cell proliferation in the spleen was determined using flow cytometry (n=3 mice per group). (c) The proportion of donor CD8^+^ T cells among the total CD8^+^ T cells in the spleen was determined using flow cytometry (n=3 mice per group). (d) Analysis of the percentage of CD44^hi^CD62L^low^ cells in the splenic CD8^+^ T cells via flow cytometry (n=3 mice per group). (e) The proportion of IFN-γ-positive cells in the spleen was determined by intracellular staining (n=3 mice per group). Statistical analysis was performed by two-way ANOVA followed by Tukey’s test (b-e). (f) Pharmacokinetic studies of AP-EVs were carried out using ^64^Cu-labeled AP-EVs with PET imaging. Representative maximum intensity projection (MIP) PET images are shown. (g) The average radioactivity concentration across tissues such as blood, urine, lung, heart, liver, kidney, spleen, intestine, muscle, fat, skin, and brown fat was presented as the percentage of injected dose per gram of tissue (%ID/g) (n=4 mice per group). (h-i) Recipient mice were treated with AP-EVs prior to OT-I T cells transfer at specific intervals: 6, 5, 4, 3, 2, or 0 days. Four days post-transfer, the proliferation of adoptively introduced OT-I T cells was quantified via flow cytometry (n=3 mice per group).

Furthermore, we conducted a pharmacokinetic analysis of AP-EVs following intravenous injection. We examined the pharmacokinetics of ^64^Cu-labeled AP-EVs using positron emission tomography (PET) imaging^28^ and found that AP-EVs predominantly accumulated in the spleen and liver (Fig.3f, g). To clarify the duration of the functional persistence of AP-EVs in vivo, we administered them several days prior to the transfer of OT-I T cells. Although a substantial fraction of AP-EVs was cleared by day 2, sufficient activating capacity remained until day 4 after AP-EV administration (Fig.3h, i).

### AP-EVs potentiate the anticancer efficiency of adoptively transferred T cells

To investigate the effector function of the AP-EVs-expanded CD8^+^ T cells in vivo, we performed an in vivo killing assay. To determine whether OT-I T cells expanded by AP-EVs in mice could specifically eliminate OVAp-coated splenocytes, we introduced a mixture of CTV-labeled OVAp-coated and CFSE-labeled non-coated splenocytes (at a ratio of 1:1) 4 days after EV administration. A marked reduction of OVAp-coated splenocytes, with their frequency decreasing from approximately 60% to 1.26%, indicating that CTLs activated by AP-EVs had high antigen-specific cytotoxicity (Fig. 4a). To evaluate the tumor-killing activity of these antigen-specific CTLs, we subcutaneously inoculated OVA-expressing EL4 cells (E.G7 cells), followed by the injection of OT-I T cells after 1 day. As depicted in Fig. 4b, the mice received three doses of AP-EVs, control EVs, or IL-2/anti-IL-2 mAb (Fig.4b, c). For initial administration, AP-EVs were loaded with pMHCI, CD80, and IL-2. For the two subsequent administrations, we used AP-EVs expressing pMHCI and IL-2, omitting CD80 to avoid stimulating the inhibitory receptor CTLA-4. As previously reported, IL-2/anti-IL-2 mAb only induced a marginal delay in cancer growth and limited the extension of survival^29^. The administration of AP-EVs significantly delayed cancer progression, and six out of nine mice completely rejected E.G7 cells. Moreover, mice that rejected E.G7 cells showed pronounced resistance upon subsequent E.G7 cell rechallenge (Fig. 4d, e). These findings demonstrate that the administration of AP-EVs stimulates the effector function of antigen-specific CTLs and facilitates the generation of antigen-specific memory CD8^+^ T cells.

**Figure 4.**
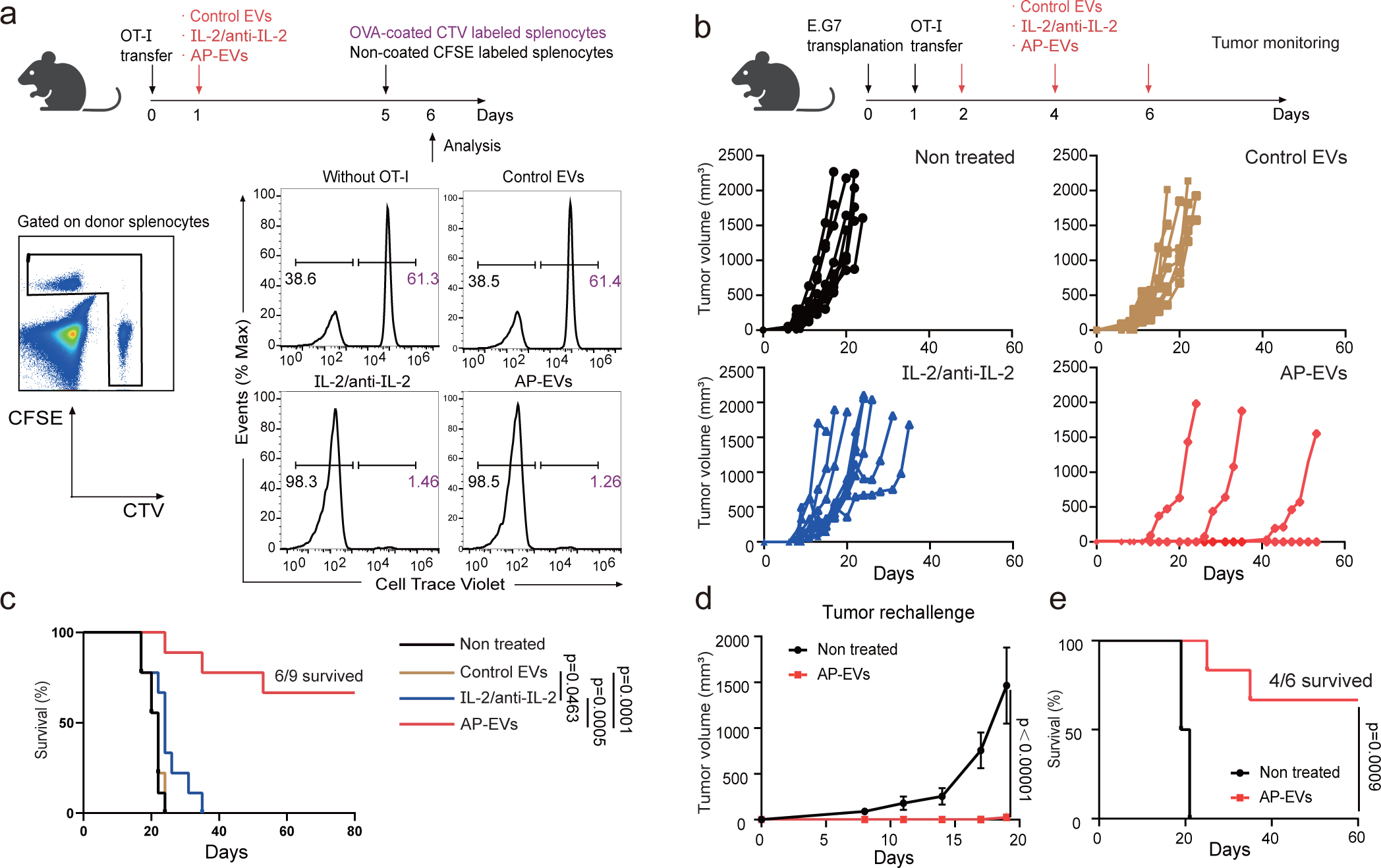
Effect of AP-EV administration on anticancer-immunity. (a) The experimental flow of in vivo killing assay with adoptively transferred OT-I T cells was presented. CD45.1^+^CD45.2^+^ recipient mice were treated with CD45.1 OT-I T cells, followed by an injection of either control EVs, IL-2/anti-IL-2 mAb, or AP-EVs. The survival of donor CTV-labeled OVAp-pulsed CD45.2^+^ splenocytes and CFSE-labeled non-pulsed CD45.2^+^ splenocytes was assessed using flow cytometry. Data show the ratio of OVAp-pulsed splenocytes (CTV-labeled) to non-pulsed splenocytes (n = 3 mice per group). (b) E.G7 cells were transferred into recipient mice. The following day, mice were treated with either control EVs, IL-2/anti-IL-2 mAb, or AP-EVs three times at a three-day interval. Tumor growth curves represent the following mouse groups: untreated(black), control EVs(brown), IL-2/anti-IL-2 mAb(blue), or AP-EVs(red) treated (n = 9 mice per group). (c) The Kaplan-Meier survival curve illustrates overall survival. Statistical analysis was performed via a Log-rank (Mantel-Cox) test. (d) Mice that previously rejected E.G7 post AP-EV treatment were later subcutaneously challenged with 1 x 10^5^ E.G7 cells. Displayed tumor growth curves compare untreated mice with those treated using AP-EVs. P-values were ascertained via a two-way ANOVA, followed by a Bonferroni test. (e) Kaplan-Meier survival curve for mice subjected to tumor rechallenge. Statistical analysis was performed using a Log-rank (Mantel-Cox) test (n = 6 mice per group).

### AP-EVs amplify endogenous antigen-specific CD8^+^ T cells and enhance anticancer immunity

Endogenous CD8^+^ T cells that recognize a specific peptide-MHC complex are present at frequencies ranging from 1 to 100 per million CD8^+^ T cells^30^. The OVA-specific endogenous CD8^+^ T cells comprise approximately 0.02% of the total CD8^+^ T cells in peripheral blood mononuclear cells (PBMCs) from untreated mice (Fig.5a). We investigated the capacity to expand this rare population in mice by administering AP-EVs, control EVs, or an IL-2/anti-IL-2 mAb on days 0, 2, and 4 and subsequently analyzed them on day 7. The AP-EVs triggered the clonal expansions of OVAp-specific CD8^+^ T cells, representing 4.24% of all CD8^+^ T cells. In contrast, control EVs or IL-2/anti-IL-2 mAb treatment failed to selectively expand OVAp-specific CD8^+^ T cells (Fig.5a). To verify the effector function of these expanded OVAp-specific CD8^+^ T cells, we performed an in vivo killing assay. We introduced a mixture of CD45.2 OVAp-coated splenocytes and CD45.1 non-coated splenocytes (at a ratio of 1:1) two days after the third EV administration.

**Figure 5.**
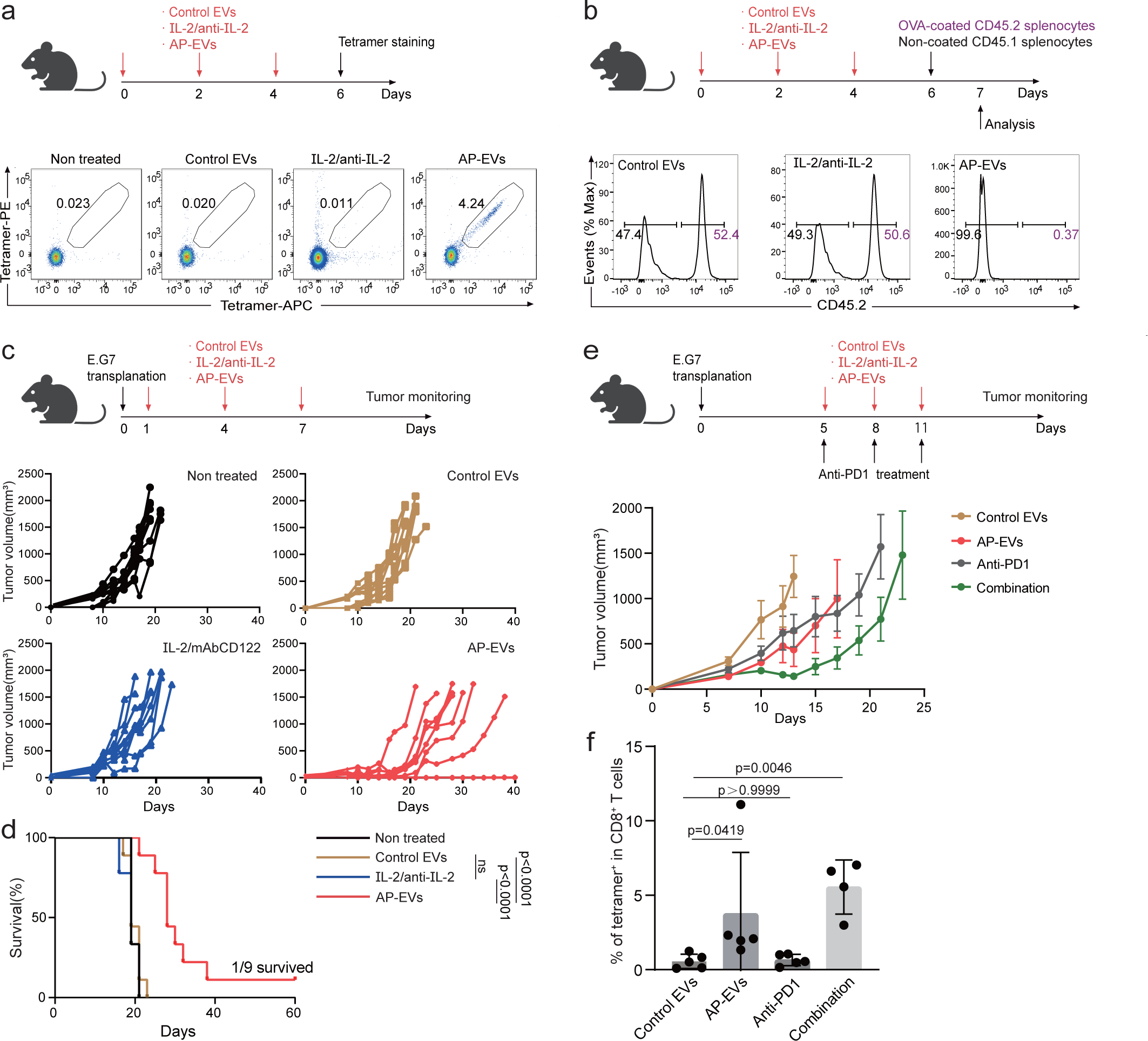
Effect of AP-EVs on the expansion of endogenous antigen-specific CD8^+^ T cells and cancer immunity. (a) OVAp-specific CD8^+^ T cells in PBMCs from mice which were treated with either control EVs, IL-2/anti-IL-2 mAb, or AP-EVs were detected by using tetramer staining on day 6. The percentages of tetramer-positive cells in untreated and control EVs, IL-2/anti-IL-2 mAb, or AP-EV treated mice are presented (n = 4 mice per group). Data are representative of two independent experiments. (b) The experimental flow of in vivo killing assay was presented. The survival rate of the donor splenocytes in recipient mice which were injected with either control EVs, IL-2/anti-IL-2 mAb, or AP-EVs was determined via flow cytometry. The ratios displayed compare OVAp-pulsed (CTV-labeled) with non-pulsed splenocytes. (c) E.G7 cells were transferred into recipient mice. The following day, mice were treated with either control EVs, IL-2/anti-IL-2 mAb, or AP-EVs three times at a three-day interval. Tumor growth curves illustrate the following groups: untreated (black), control EVs (brown), IL-2/anti-IL-2 mAb (blue), and AP-EVs (red) treated (n=9 mice per group). (d) Kaplan-Meier survival curve depicting overall survival rates. Statistical analysis was performed via a Log-rank (Mantel-Cox) test. (e) E.G7 tumor-bearing mice were treated three times, at three-day intervals starting from day 5, with one of the following: control EVs, AP-EVs, anti-PD1 antibody, or a combination of AP-EVs and anti-PD1 antibody. Tumor growth curves corresponding to each treatment group: control EVs, AP-EVs, anti-PD1 antibody, or AP-EVs with anti-PD1 antibody. During the tumor measurement process, one mouse in a combination therapy group unexpectedly died despite having a small tumor size. Data from this mouse were excluded from subsequent analyses. Data are shown as the standard error of the mean. (f) OVAp-specific CD8^+^ T cells in PBMCs were detected on day 14 using tetramer staining. Percentages of tetramer-positive cells from mice treated with control EVs, AP-EVs, anti-PD1 antibody, or AP-EVs with anti-PD1 antibody are presented (n=5 mice per group). P values were determined via using one-way ANOVA, followed by Dunn’s multiple comparisons tests.

OVAp-coated splenocytes were eliminated from mice that received AP-EVs (Fig.5b). This effect was not observed in the mice treated with control EVs or IL-2/anti-IL-2 mAb. Furthermore, we investigated whether AP-EVs exert anticancer effects via endogenous CTLs. We subcutaneously injected E.G7 cells into C57BL/6 mice and administered AP-EVs, control EVs, or IL-2/anti-IL-2 mAb. Notably, AP-EV2 treatment resulted in a substantial delay in cancer progression and an extension of survival compared to that in the control groups (Fig.5c, d). Intriguingly, AP-EVs accumulated at the tumor site, implying the potential activation of tumor-infiltrating lymphocytes by AP-EVs (Extended Data Fig.6a, b, c). Next, we explored the efficacy of AP-EVs against established tumors. E.G7 cells were subcutaneously injected into mice and allowed to proliferate until they reached a size of 100 mm^3^. The mice were administrated AP-EVs, anti-PD-1 antibodies, or a combination of both. Monotherapy using either AP-EVs or anti-PD-1 antibody resulted in only a slight delay in tumor progression, whereas combination therapy markedly inhibited tumor growth (Fig.5e). Notably, we observed an increased percentage of antigen-specific CD8^+^ T cells among the total CD8^+^ T cells in the PBMCs of mice treated with combination therapy, reflecting a synergistic effect between AP-EVs and the anti-PD-1 antibody (Fig.5f). These findings highlight the effectiveness of AP-EVs not only in the selective expansion of endogenous antigen-specific CD8^+^ T cells but also in enhancing anticancer immunity, particularly when employed with an anti-PD-1 immune checkpoint inhibitor.

## Discussion

Extracellular vesicles, including exosomes, have attracted significant attention owing to their potential as drug delivery systems, diagnostic tools, agents for tissue regenerative medicines, and immune response modulators. Unlike cell-based therapies, EVs offer several advantages, including reduced vulnerability to microvascular entrapment, minimal risk of malignant transformation, and superior stability compared to therapeutic cells. Hence, there has been considerable interest in developing engineered EVs capable of activating immune responses for use in cancer immunotherapy in clinical settings. However, their efficacy has been limited^31^.

To overcome these limitations, we engineered AP-EVs equipped with multiple immune modulators on a single-EV platform. This design allows targeted T cells to concurrently receive various signals, such as TCR, costimulatory, and cytokine signals. A key advantage of AP-EVs is their adaptability to modifications, which facilitates the incorporation of a diverse array of cytokines and costimulatory signals. In this study, we integrated IL-2 and CD80 as prototypical cytokines and costimulatory molecules, respectively, into EVs. However, the use of alternative cytokines and signals, such as IL-12 or 4-1BBL, is conceivable. For instance, EVs containing pMHCII, CD80, and IL-12 are anticipated to promote antigen-specific Th1 differentiation, while EVs embedded with pMHCII, IL-2, and TGF-β may induce antigen-specific Tregs, presenting potential therapeutic avenues for autoimmunity. Additionally, immune checkpoint inhibitors can be applied to AP-EVs to generate anti-PD-1 scFV-tetraspanin fusion proteins^32^. This adaptability in selecting costimulatory molecules, cytokines, and other immune modulators provides flexibility to customize AP-EVs, making them suitable for various conditions, including cancer and autoimmune diseases.

Dendritic cell-based cancer immunotherapies, particularly DC vaccines, have emerged as promising therapeutic strategies against malignancies such as acute myeloid leukemia, myelodysplastic neoplasms, and various solid tumors^33,34^. Their ability to present tumor-specific antigens and stimulate robust antitumor responses has increased interest in their clinical use. While many studies have shown that DC vaccines are safe and feasible, several limitations persist, including the modest cellular immune response they elicit, high associated costs and intricate production processes^35^. In contrast, owing to their scalable production, ease of modification, and inherent stability stemming from their non-cellular composition, AP-EVs may be a more efficient strategy than DCs vaccines. Furthermore, the adaptability of AP-EVs allows the integration of key insights from DC vaccine research, such as specific antigen selection, methods for more robust immune responses, and safety protocols. By leveraging these strategies within the AP-EV design, we can potentially enhance the effectiveness of cancer immunotherapy by combining the strengths of both approaches.

Several modified cytokines have been developed to boost their functions while avoiding the adverse effects of systemic administration. For examples, engineered IL-2 variants have been developed to mitigate the side effects of IL-2 by selectively activating CD8^+^ memory T and NK cells^18,36^. However, these IL-2 variants lack antigen specificity and cause expansion of the entire repertoire of CD8^+^ T cells. Orthogonal pairs of IL-2/IL-2R offer a way to specifically target T cells^37^. However, this method requires genetic modification of T cells and may inadvertently activate the immune system through repetitive injections, because orthogonal pairs of IL-2/IL-2R are not recognized as self-antigens. The AP-EVs technology circumvents these challenges by allowing the selective delivery of IL-2 to antigen-specific T cells, owing to the simultaneous expression of pMHCI and IL-2 on EVs. This feature could lead to a safer and more effective activation of antigen-specific CD8^+^ T cells.

## Conclusions

In conclusion, AP-EVs, which express several immune modulators along with pMHCI, have the potential to selectively deliver immune modulators to antigen-specific T cells, thereby introducing a novel approach to cancer immunotherapy. While our study demonstrates the potential of AP-EVs, further research is needed to optimize their design and evaluate their safety and efficacy in non-clinical and clinical trials. Overall, AP-EVs are expected to facilitate the development of novel immunotherapies.

## Methods

### Cell lines

Human embryonic kidney cells (HEK293 or HEK293T) and cells of a retroviral packaging cell line (PLAT-A) were cultured in Dulbecco’s Modified Eagle Medium (DMEM; Thermo Fisher Scientific) supplemented with 10% heat-inactivated fetal calf serum (FCS; Thermo Fisher Scientific), 100 U/ml penicillin, and 100 U/ml streptomycin (FUJIFILM Wako). Cells of an ovalbumin-expressing murine lymphoma cell line, an EL4 derivative (E.G7; ATCC, CRL-2113), and murine lymphocytes were cultured in RPMI (Nacalai Tesque) supplemented with 10% FCS, 1X of non-essential amino acid (Nacalai Tesque), 1 nmol/L sodium pyruvate (Nacalai Tesque), 100 U/ml penicillin, 100 U/ml streptomycin (FUJIFILM Wako), and 0.05 μM 2-Mercaptoethanol (Thermo Fisher Scientific). All the cells were cultured at 37 °C in a humidified atmosphere containing 5% CO_2_. To prepare EV-free FCS, FCS was mixed with 50% PEG-10,000 (Merck) at a 5:1 ratio and rotated at 4 °C for 3 h; next, PEG was removed by centrifuging at 2,000 g for 20 min and the supernatant was filtered through a 0.22-μm strainer and used as EV-depleted FCS. The AP-EV-producing cell line was cultured in advanced DMEM supplemented with 2% EV-depleted FCS, 1 nmol/L sodium pyruvate, 100 U/ml penicillin, and 100 U/ml streptomycin.

### Mice

C57BL/6 and CD1 mice were purchased from Japan SLC and Jackson laboratory, respectively. OT-I TCR transgenic mice (CD45.1) have been previously described^38^. All mice were housed in a specific pathogen-free facility. All animal experiments were performed following a protocol approved by Kanazawa University.

### Preparation and purification of AP-EVs

HEK293 cells stably expressing OVA peptide-single chain trimer-CD81-IL2 and CD80-CD9 were generated using retroviral systems. The cell culture supernatant was first centrifuged at 300 g for 5 min to remove cell debris and then re-centrifuged at 1,200 g for 20 min to remove cell debris and apoptotic bodies. Next, the supernatant was again centrifuged at 10,000 g for 30 min to remove apoptotic bodies and large extracellular vesicles. AP-EVs were isolated from the supernatant through centrifugation at 100,000 g for 4 h; the pellet obtained was washed with PBS and the AP-EV concentration was determined through a BCA assay (Thermo Fisher Scientific).

### Nanoparticle Tracking Analysis (NTA)

The number and characteristics of AP-EVs were determined using the NanoSight LM10 (Malvern Panalytical, Malvern, United Kingdom). In brief, 600 μL of a diluted AP-EVs solution was placed on a sample stage and the motion of AP-EVs was captured at a camera level of 15 for 30 s. Three distinct fields were recorded and data were analyzed using the NTA3.1 software with the application of a detection threshold of 3.

### Confocal microscope imaging

EVs were isolated from CD81-GFP- and CD9-RFP-co-transfected HEK293T cells and a single EV was analyzed by confocal fluorescence microscopy (Nikon A1R). Fluorescence stacks (GFP & mCherry Chroma filter-set; 5 μm; 130 nm steps) were acquired, deconvolved, and processed into maximum intensity projection images using the MIP-Delta vision software. A colocalization channel with an intensity threshold of 300 au was generated by Imaris9.3. Spots in the green, red, and colocalization channels were quantified using the Imaris software, with the application of a minimum diameter and quality threshold of 0.2 μm and 500 au (intensity at the center of the spot), respectively.

### Antibodies

Antibody staining was performed according to standard procedures. Monoclonal antibodies for surface staining, including antibodies against CD3 (clone:17A2), CD4 (clone: GK1.5), CD8α (clone:53-6.7), CD19 (clone:6D5), CD25 (clone: PC61), NK1.1 (clone: S17016D), mTCR-Vβ5.1,5.2 (clone: MR9-4), CD45.1 (clone: A20), CD45.2 (clone:104), CD44 (clone: IM7), CD62L (clone: MEL-14), H-2K^b^ bound to SIINFEKL-APC (clone:25-D1.16), CD80 (clone:16-10A1), and IL-2 (clone: JES6-5H4), were purchased from BioLegend. Intracellular staining was performed using antibodies against IFN-γ (clone: XMG1.2), granzyme B (clone: MF-14), and Foxp3 (clone:53-6.7) and the True-Nuclear Transcription Factor Buffer set (BioLegend). OVA-specific CD8^+^ T cells were quantified using the H-2K^b^/OVA _(257-264)_ tetramer (Tetramer shop) according to the instructions of the manufacturer.

### Flow cytometric analysis of EVs

The surface expression levels of pMHCI, CD80, and IL-2 on EVs were assessed using the PS Capture Exosome flow cytometry kit (FUJIFILM Wako) according to the instructions of the manufacturer. In brief, the cell culture supernatant was centrifuged at 10,000 g and incubated with EVs capture beads for 1 h. Next, EVs-bound beads were stained with fluorescence-labeled antibodies. Finally, after washing the stained EV-bound beads, they were analyzed using a CytoFLEX flow cytometer (Beckman Coulter) and the resulting data were processed using the FlowJo v.10.4.1 software.

### AFM analysis

EVs were immobilized on a poly-L-lysine (PLL)-modified mica substrate. A Bruker BioScope Resolve AFM system operated in the Peak Force Tapping mode was used for imaging and nanomechanical analysis following the method described by Yurstver et al.^39^ AFM images were captured using 240 AC-NG cantilevers with the spring constant set using the thermal tuning method. The peak force tapping amplitude, frequency, and setpoint were set to 20–50 nm, 1 kHz, and 0.5–2 nN, respectively. For nanomechanical analysis, the Young’s modulus was measured using quantitative nanomechanical mapping (QNM) and the Sneddon contact mechanical model, scanning a typical 256 x 256-pixel area. AFM image rendering and nanomechanical data processing were performed using the NanoScope Analysis software version 1.9.

### IL-2 bioassay

CTLL-2 cells were seeded in 96-well plates at a density of 8 × 10^4^ cells/well and serial AP-EVs dilutions were added to each well. Cell viability was measured after 3 days of culturing through the WTS-I colorimetric assay (Nacalai Tesque).

### In vitro T cell proliferation assay

LN T cells from OT-I transgenic mice were labeled with 1 μM CTV (Thermo Fisher Scientific) at 37 ℃ for 3 min. A total of 2 × 10^5^ CTV-labeled OT-I T cells were co-cultured with AP-EVs or control EVs at concentrations of 3, 1, 0.3, and 0.1 μg/ml, or with anti-mouse CD3/CD28 beads (Thermo Fisher Scientific) for 3 days. OT-I T cell proliferation was evaluated by flow cytometry.

### Adoptive T cell transfer

LN T cells from OT-I (CD45.1) and C57BL/6 mice (CD45.2) were mixed at a 1:1 ratio and labeled with CTV. A total of 2 × 10^6^ T cells were intravenously injected into CD45.1/CD45.2 recipient mice. One day after, the recipient mice were treated with either 50 μg of AP-EVs or control EVs, or IL-2/IL-2mAb, which comprised 1.5 μg of mouse recombinant IL-2 (Biolegend) and 50 μg of anti-IL-2 monoclonal antibodies (S4B6, Bio X cell). After 4 days, the spleens and LNs of the mice were collected and single-cell suspensions were prepared and analyzed by flow cytometry.

### Toxicity testing

AP-EVs were administered to CD1 mice at a dose of 1.25 mg/Kg. Mouse body weights were monitored on days 1, 3, 7, and 14. On days 1 and 14, the mice were euthanized and subjected to hematological, blood chemistry, and histopathological examinations.

### Pharmacokinetics of EVs

EVs were labeled with ^64^Cu following a previously described method^28^. PET analysis was performed using a Focus 220 MicroPET scanner (Siemens Co., Knoxville, TN, USA). ^64^Cu-labeled EVs were intravenously administered to anesthetized recipient mice at the start of the PET scan. PET images were reconstructed using MicroPET Manager (v. 2.4.1.1; Siemens Co.). The radioactivity of each pixel was decay-corrected from the time of injection and expressed as the standardized uptake value (SUV), which was calculated using the formula, SUV = (tissue radioactivity concentration [MBq/cm3])/(injected radioactivity [MBq]/body weight [g]).

### In vivo killing assay

Recipient mice were administrated either 200 μg of AP-EVs or control EVs, or IL-2/IL-2 mAb. As shown in the protocol depicted in Fig. 4, four days following EV administration, the mice were administered target splenocytes (2 x 10^6^) which comprised an equal mix of OVAp-coated CTV-labeled splenocytes and non-coated CFSE-labeled splenocytes. Furthermore, as shown in the protocol depicted in Fig. 5, two days after the third EV administration, mice were administered an equal-ratio mixture of CD45.2 OVAp-coated splenocytes (1 x 10^6^) and CD45.1 non-coated splenocytes (1 x 10^6^). For both protocols, mouse spleens and lymph nodes were harvested 20 h following splenocyte administration. Single-cell suspensions were prepared from these organs and analyzed by flow cytometry.

### Tumor model

To establish the E.G7 tumor model, a total of 1 x 10^5^ E.G7 cells were subcutaneously inoculated into C57BL/6 mice. On the following day, tumor-bearing mice were intravenously administered 1 × 10^6^ CD45.1 OT-I T-cells. This was followed by the administration of 200 μg of AP-EVs, control EVs, or IL-2/IL-2 mAb every three days starting from day 2, for a total of three administrations. The inoculation procedure was similar to that used for the tumor model established without adoptive OT-I T cell transfer; however, treatment was initiated on day 1 post E.G7 cell inoculation. Tumor volumes were measured every 2–3 days using a digital caliper and were calculated using the formula, length (mm) × width^2^ (mm^2^) × 0.5. Prior to the anti-PD1 combination therapy, E.G7 cells were subcutaneously injected into the mice. Treatment was initiated when the tumors were 50–100 mm^3^ in size. Then, the mice were randomly allocated to one of four groups i.e., the AP-EVs, control EVs, 200 μg anti-PD1 antibody, and combination therapy groups. The 200 μg AP-EV/control EV treatments were administered every two days for a total of three administrations. The anti-PD1 treatment was administered every three days for a total of three administrations. Tumor volume measurements and calculations were performed as described above.

## ACKNOWLEDGMENTS

We thank T. Yoshida for the helpful discussions and Dr. Y. Watanabe, Dr. Y. Wada, and Ms. E. Hayashinaka from RIKEN for their assistance in reconstructing the PET images. This work was supported by the Japan Science and Technology Agency (JST) Precursory Research for Embryonic Science and Technology (PRESTO) (No. JPMJPR19HA to TY), Science and Technology Platform for Advanced Biological Medicine from the Japan Agency for Medical Research and Development (AMED) (No. 22am0401019h0004 to RH), Core Research for Evolutional Science and Technology (CREST) (No. JPMJCR18H4 to RH), and AMED (No. JP23ak0101178 to HM). TY is supported by The Kanae Foundation for the Promotion of Medical Science and MSD Life Science Foundation, Public Interest Incorporated Foundation.

## Author Contributions

TY and RH were responsible for the conception and experimental strategy of the study. XL TY, SI, TVL, and MU designed and analyzed the experiments. DB performed the AFM analysis. SW and HM analyzed the Pharmacokinetics of EVs. SH, KM, and TN analyzed the toxicity of EVs.

