## Extended Data Fig. 1-6 for "Surface-engineered extracellular vesicles to modulate antigen-specific T cell expansion for cancer immunotherapy"

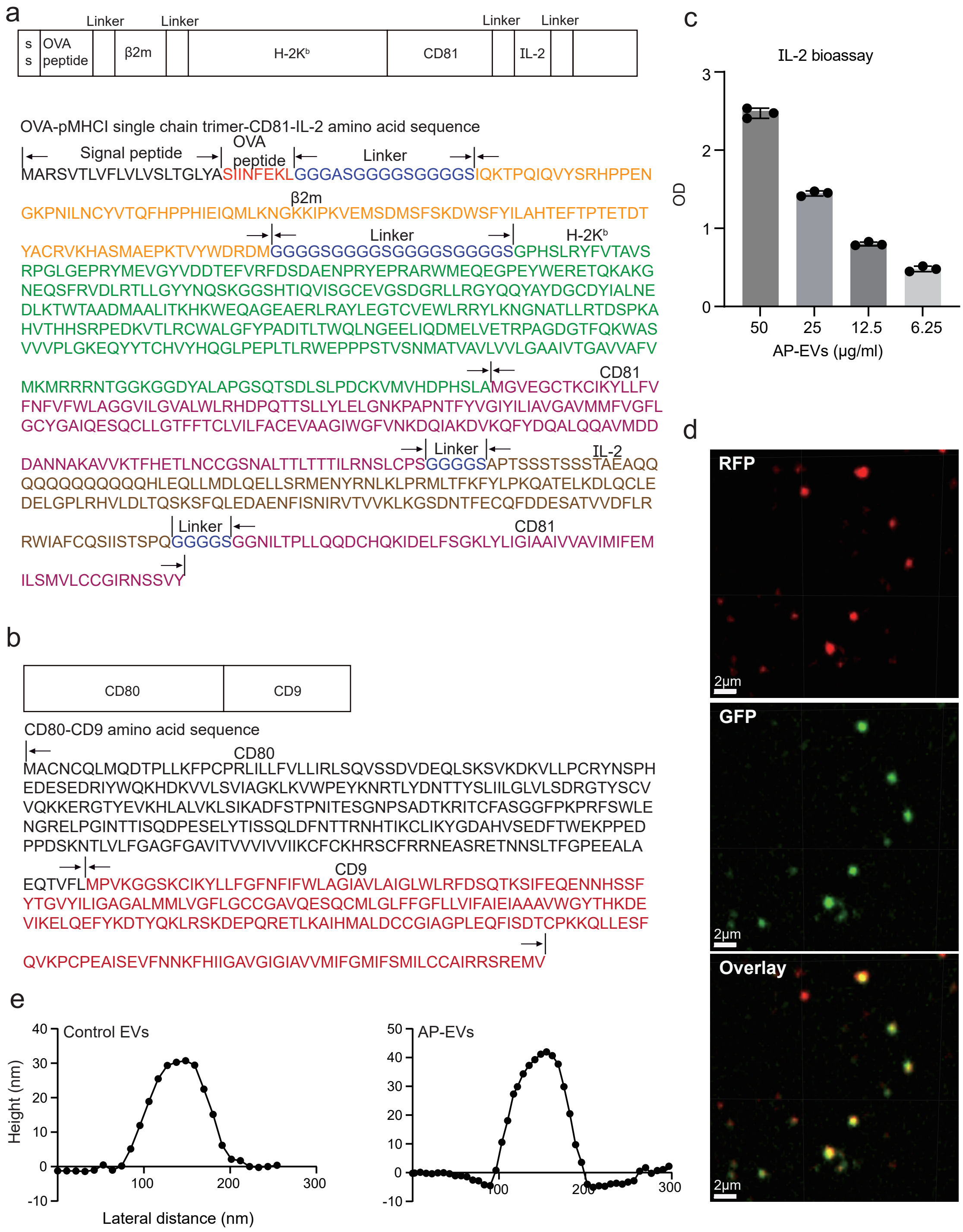


**Extended Data Fig. 1 | Characteristics of antigen presenting EVs (AP-EVs).**
**(a)** Schematic illustration of protein fusion. Amino acid sequence of the OVA-pMHCI single chain trimer-CD81-IL-2. The OVA-pMHCI single chain trimer was fused with CD81 and IL-2. **(b)** Schematic illustration of protein fusion. Amino acid sequence of CD80-CD9 fusion proteins. CD80 was conjugated with the C terminus of CD9. **(c)** CTLL-2 cells were cultured in the presence of serial AP-EV dilutions. After 3 days of culturing, cell viability was assessed through the WST-I assay. Absorbance was determined using a spectrophotometer at 450 nm. **(d)** Confocal microscopic analysis of EVs expressing CD81-RFP and CD9-GFP; scale bar: 2 μm. **(e)** Size distribution and height of isolated EVs as determined by atomic force microscopy (AFM). Data (c, d, e) are representative of the results of two independent experiments.

**Extended Data Figure 1 Lyu et al.**

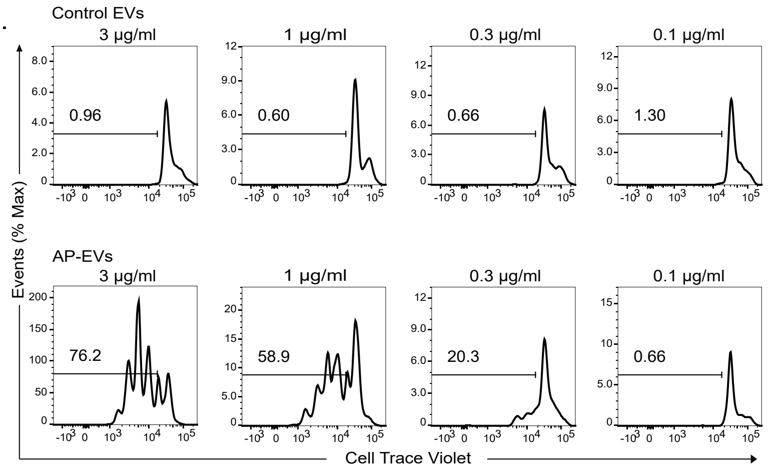


**Extended Data Fig. 2 | AP-EVs selectively expand antigen-specific CD8$^+$ T cells in vitro.** CTV-labeled OT-I T cells were cultured with serial dilutions of control EVs or AP-EVs for three days. Proliferated OT-I T cells were analyzed by flow cytometry. Data are representative of the results of two independent experiments.

**Extended Data Figure 2 Lyu et al.**

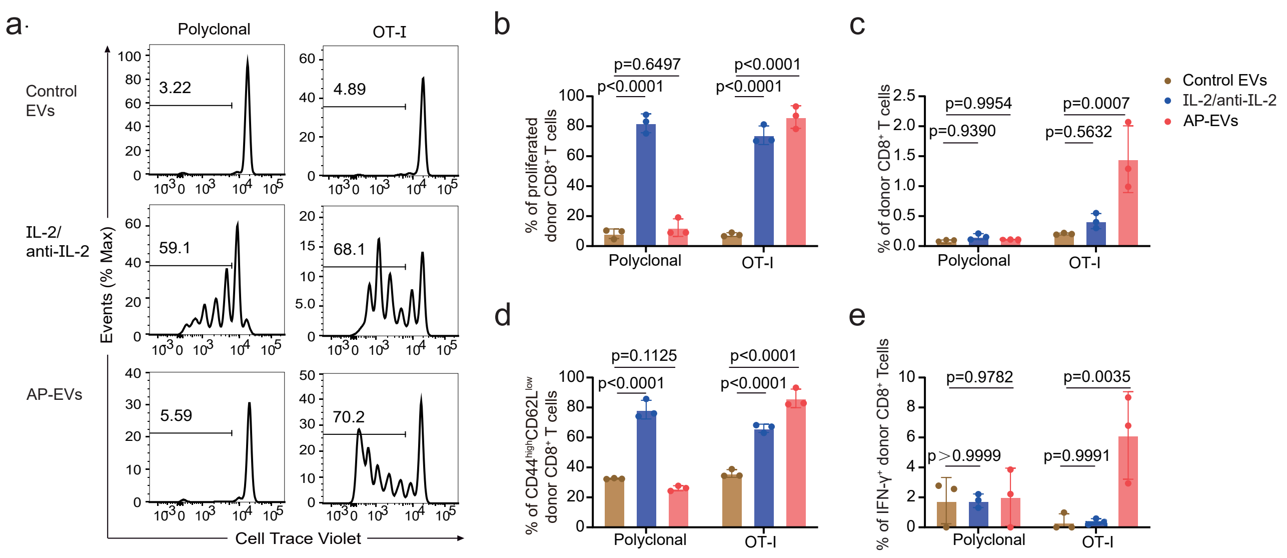


**Extended Data Fig. 3 | Activation of antigen-specific CD8⁺ T cells by AP-EVs in lymph nodes.** **(a)** CTV-labeled OT-I T cells (CD45.1) and polyclonal T cells (CD45.2) were co-administered to recipient mice (CD45.1⁺ CD45.2⁺). One day after administration, the mice received control EVs, the IL-2/anti-IL-2 mAb, or AP-EVs. Mouse LNs were harvested three days following AP-EV administration and T cell proliferation was assessed by flow cytometry (n = 3 mice per group). **(b)** T cell proliferation percentage in mouse LNs. **(c)** Percentage of donor-derived CD8⁺ T cells in mouse LNs. **(d)** Percentage of CD44$^{high}$ CD62L$^{low}$ donor CD8⁺ T cells in mouse LNs. **(e)** Percentage of IFN-γ⁺ cells in mouse LNs. Statistical analysis was performed by two-way ANOVA followed by Tukey's test.

**Extended Data Figure 3 Lyu et al.**

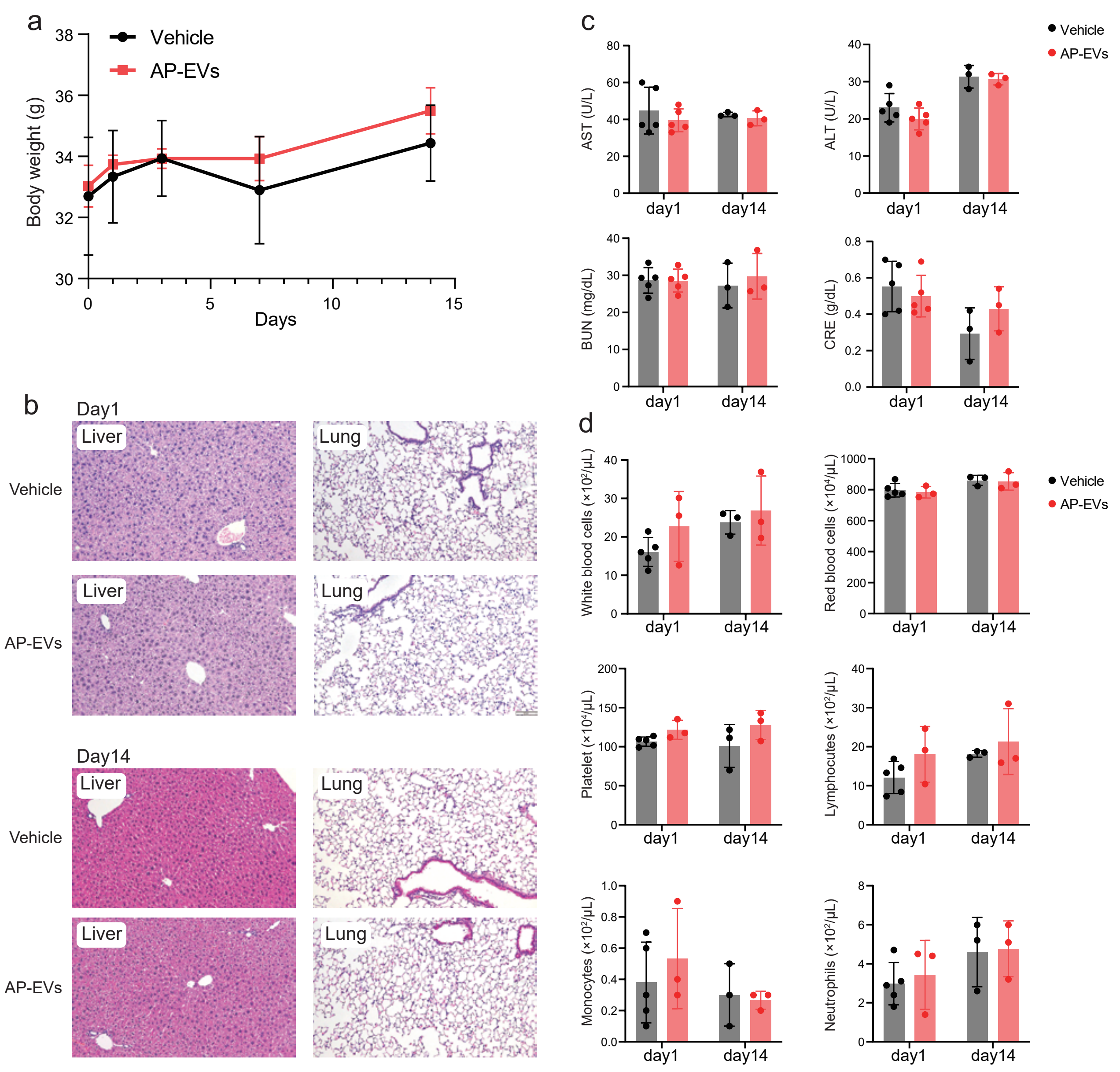


**Extended Data Fig. 4 | No adverse effects were observed following AP-EV administration.** Mice were intravenously administered 1.25 mg/kg of AP-EVs or vehicle (n = 3-5 mice per group). **(a)** Mouse body weight was measured following treatment with AP-EVs or vehicle for 0, 1, 3, 7, and 14 days. **(b)** Histopathological examination of liver and lung tissues following treatment for 1 and 14 days. **(c)** Blood chemistry analysis at days 1 and 14 post-treatment. **(d)** Hematological assessment at days 1 and 14 post-treatment.

**Extended Data Figure 4 Lyu et al.**

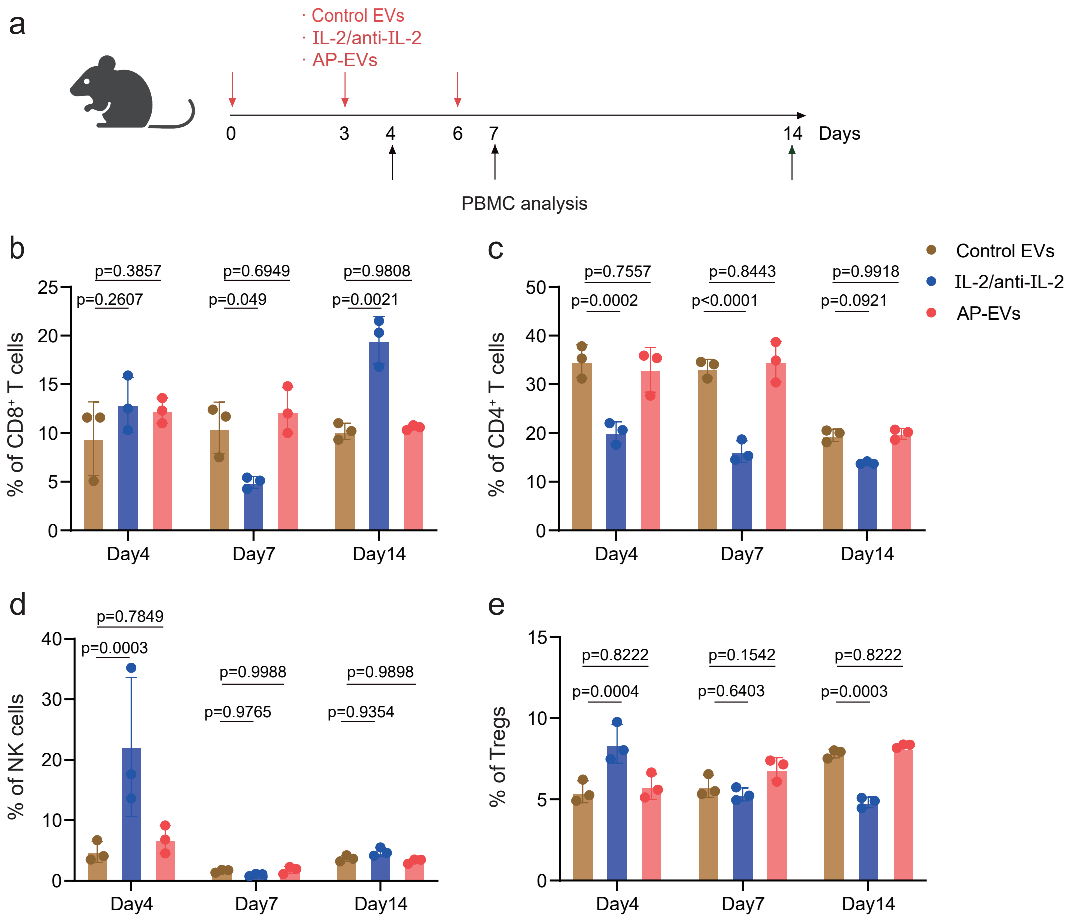


**Extended Data Fig. 5 | AP-EV administration does not expand NK or regulatory T cells.**
**(a)** Schematic representation of the AP-EV administration protocol. C57BL/6 mice received either 50 μg of control EVs, IL-2/anti-IL-2 mAb, or AP-EVs, administered thrice on days 0, 3, and 6. PBMCs were harvested on days 4, 7, and 14 to analyze the effects of the treatments on CD4⁺ T cell, CD8⁺ T cell, NK cell, and Tregs populations (n = 3 mice per group). **(b)** Blood CD8⁺ T cell percentage on days 4, 7, and 14. **(c)** Blood CD4⁺ T cell percentage on days 4, 7, and 14. **(d)** Blood NK cell percentage on days 4, 7, and 14. **(e)** Blood Tregs cell percentage on days 4, 7, and 14. Statistical analysis was performed by two-way ANOVA followed by Tukey's test.

**Extended Data Figure 5 Lyu et al.**

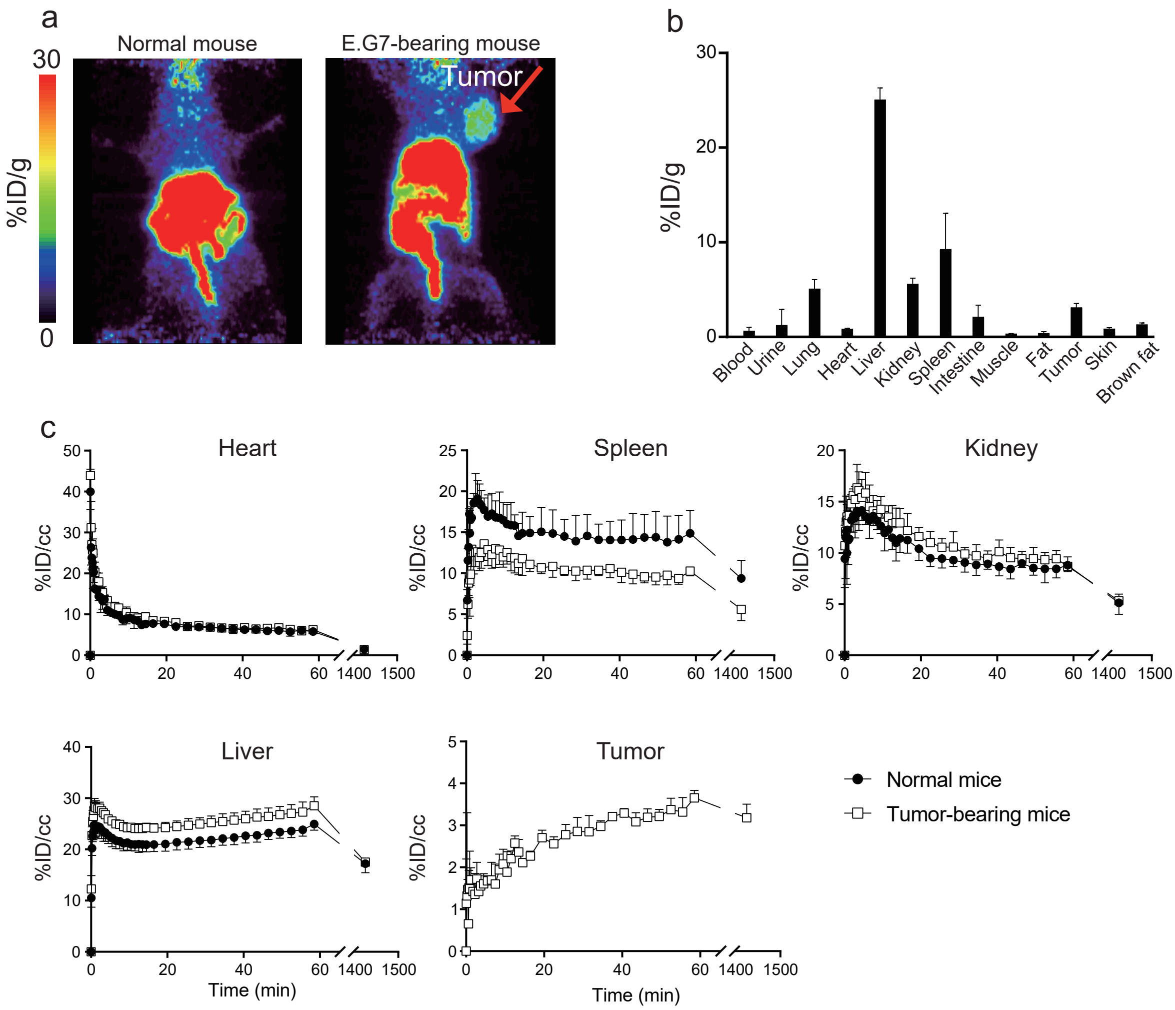


**Extended Data Fig. 6 | AP-EV tumor site accumulation.**

**(a)** AP-EVs were labeled with $^{64}$Cu and administered to untreated C57BL/6 mice and mice bearing E.G7 tumors. AP-EV accumulation was evaluated 24 h following administration. Representative maximum intensity projection PET images are presented (n=4 mice per group). **(b)** The mean radioactivity levels of $^{64}$Cu-labeled AP-EVs in various samples, including blood, urine, lung, heart, liver, kidney, spleen, intestinal, muscle, fat, skin, tumor, and brown fat samples, are expressed as the percentage of the injected dose per gram of sample (%ID/g). **(c)** Time-activity curve present the percentage of the injected dose per cubic centimetre of tissue (%ID/cc), including heart, spleen, kidney, liver, and tumor tissues.

**Extended Data Figure 6 Lyu et al.**
